# Condition Medium of Glioblastoma Cell Lines Decreases the Viability of Glioblastoma Cells by Modulating Gene Expression Profile

**DOI:** 10.1101/2021.09.11.459916

**Authors:** Mervenur Yavuz, Sıddıka Akgül, Egemen Kaya, Turan Demircan

## Abstract

Grade IV neoplasm of the central nervous system, GBM, is associated with poor prognosis and relatively short overall survival. Due to the current limitations in treatment methods, GBM is characterized as an incurable disease, and research to advance therapeutic options is required. Conditioned medium is commonly used in *in-vitro* studies complementary to animal experiments to simulate tumor microenvironment and has the potential to challenge and expand our current understanding of secretome effect on tumor characteristics. This study aimed to investigate the effects of conditioned mediums of GBM cell lines on each other. Conditioned mediums’ cellular and molecular effects were evaluated using commonly employed techniques such as MTT assay, colony formation assay, wound healing assay, EdU labeling-based flow cytometry, and qRT-PCR. Our study demonstrated that conditioned medium harvested from U87 or LN229 cells at 48^th^ h exhibited an anti-growth activity on each other by changing the gene expression pattern. Furthermore, the conditioned medium of LN229 decreased the migration capacity of U87 cells, and the conditioned medium of U87 cells significantly suppressed the LN229 proliferation. We believe that this initial work provides new insights for a better understanding of GBM cell lines’ secretome roles and highlights the necessity of further studies to unveil the secretome content.

**Highlights:**

- Conditioned medium harvested from GBM cells at different time points displayed various effects.
- Conditioned medium of GBM cell lines harvested at 48^th^ h decreased the viability of each other.
- The expression level of anti-and pro-proliferative genes is altered upon condition medium treatment.

## 1. Introduction

Glioblastoma multiforme (GBM), grade IV astrocytoma, is the most aggressive and lethal form of central nervous system cancer that arises from the transformation of glial cells. The incidence of GBM increases with age and differs by sex (Omuro and DeAngelis, 2013; Stoyanov et al., 2018). GBM is classified as primary and secondary glioblastoma based on specific genetic mutations, epigenetic alterations, progressions, and origins (Ohgaki and Kleihues, 2013). Primary GBM developed in the elderly is characterized by genetic mutations of EGFR amplification, tumor suppressor PTEN gene mutation, and complete loss of chromosome 10 associated with cell proliferation, angiogenesis, migration, invasion, adhesion, and inhibition of apoptosis (Omuro and DeAngelis, 2013). Secondary GBM originates from low-grade diffuse astrocytoma, or anaplastic astrocytoma, affecting young patients with a significantly better prognosis (Ohgaki and Kleihues, 2013). TP53 tumor suppressor mutation, IDH1/2 mutations, and loss of chromosome 19q are common genetic alterations observed in secondary GBM (Ohgaki and Kleihues, 2013). Current therapy methods for glioblastomas are total resection of the tumor, chemo- and radiotherapy. Diffuse gliomas show highly invasive features; thus, curative surgical resection fails. Due to the limited benefit of current treatments, poor prognosis and recurrence of disease are the typical outcomes for GBM. Moreover, neurotoxicity caused by chemotherapy possibly triggers seconder glioblastoma formation (Davis, 2016).

The tumor microenvironment, a specialized niche composed of tumor cells, non-cancerous stromal cells, infiltrating immune cells, and cancer-associated fibroblasts (CAFs), has a remarkable effect on cancer progression and metastasis through crosstalk of secreted signaling molecules and pathways (Murgai et al., 2015). Communication with distant cells a crucial for metastasis by modulating the target site via secretome (Papaleo et al., 2017). Multiple studies have identified the roles of cancer cell-secreted molecules, including growth factors, glycoproteins, cytokines, enzymes, chaperons, signal ligands, and tumor-derived exosomes in cancer progression (Abd Elmageed et al., 2014; Bhat et al., 2021; Melo et al., 2014). Cancer cells exploit the non-cancerous cells *via* the secretome to facilitate proliferation, evasion of apoptosis, angiogenesis, and invasion (Murgai et al., 2015). The content of secreted molecules within the same tumor tissue may differ from one cell to another. Dynamic secretome seems to modify epigenetic alterations, energy metabolism, protein profile, proliferation rate, resistance to therapy, and composition of the tumor microenvironment (Farahmand et al., 2018; Guo et al., 2017; Zhuang et al., 2018). Hence, more studies to explore the content and effect of cancer secretome are required to reveal the putative pro-and anti-tumorigenic impact of the secretome.

To study GBM using the established cell culture methods in a controlled environment allows scientists to disclose the effect of conditioned medium (CM) on tumor biology (Papaleo et al., 2017). Commonly used GBM cell lines such as LN229, U87, T98G, LN18, U251 have a diverse genetic background that affects their growth, proliferation rate, migration, invasion, apoptotic status, and response to applied therapies. This study aimed to investigate the CM’s effect of GBM cell lines, LN229 and U87, on each other. Based on the obtained findings, we expect that this work will be beneficial for a more detailed exploration of the secretome content of GBM cells.

## 2. Material and Methods

### 2.1 Cell culture maintenance

Human likely glioblastoma cell lines LN229 and U87, and human astroglial cell line SVG p12 (American Type Culture Collection #Catalog numbers: CRL-2611^™^, HTB-14, CRL-8621; respectively), were used in this study. U87 and SVG p12 cell lines were cultured in 10% heat-inactivated fetal bovine serum (FBS; Gibco, Grand Island) and 1% penicillin/streptomycin (Pen/Strep; Gibco, Grand Island, USA) in high-glucose Dulbecco’s modified eagle medium (DMEM; Sigma, #Cat no: D6429). The LN229 cell line was cultured in DMEM supplemented with 5% FBS and 1% Pen/Strep. All cell lines were maintained in a humidified chamber with 5% CO_2_ at 37°C, culture mediums were replaced every other day, and cells were passaged at around 80% confluency. For the following steps, all cell lines were harvested with 0.25% Trypsin-EDTA (Gibco, #Cat no:25200056) and counted with a thoma cell counting chamber (ISOLAB, #Cat no: I.075.03.002.001) using 0.4% Trypan Blue Solution (Gibco, #Cat no: 15250661). All assays were performed in three replicates for each time point (6^th^, 12^th^, 24^th^ and 48^th^ hour CM) and control (DMEM).

### 2.2 Harvesting conditioned mediums

To collect the CMs, 0.3×10^6^ cells/well were seeded to a 6-well plate (ThermoFisher Scientific) in a 2 mL culture medium and incubated for 24 hours. The culture medium was replaced after 24 hours incubation, and CMs were collected 6, 12, and 24 hours after medium replacement. For the 48^th^ h CM, cells were seeded at 0.2×10^6^ confluency, incubated for 24 hours, the culture medium was replaced, and conditioned mediums were collected 48 hours after medium replacement. Conditioned mediums were filtered using a 0.22 μm filter (ISOLAB, #Cat no: 094.07.001) and stored at 4°C until the following experiments.

### 2.3 MTT assay

To determine the toxicity of CMs on each cell line, we performed MTT assay using CellTiter 96^®^ Non-Radioactive Cell Proliferation Assay kit (Promega, #Cat no: G400, USA). 0.1×10^5^ cells were seeded to a 96-well plate (ThermoFisher Scientific) in 100 μL medium. Cells were incubated for 24 hours for the attachment of cells, and following the 24 hours incubation, the culture medium was replaced. Each cell line was treated with other cell lines’ CMs collected at different time points. As a negative control, DMEM, and as a positive control, 10% DMSO was used. Regarding the manufacturer’s protocol, 20 μL tetrazolium dye (MTT) supplied within the kit was added to wells, and cells were incubated for four hours in a humidified chamber. Then, 100 μL solubilization solution/stop mix was added into wells to stop formazan crystal formation. Cells were incubated overnight; the formed formazan crystals were detected using SpectraMax^®^i3 (Molecular Devices) at 570 nm.

### 2.4 Colony formation assay (CFA)

CFA was conducted to determine the impact of CMs on colony-forming ability as in a previously described protocol (Demircan et al., 2021). Briefly, cells were cultured in a 96-well plate with 2×10^3^ seeding density in 100 μL medium and incubated for 24 hours. Afterward, the culture medium was replaced with the CMs or DMEM, and cells were incubated until the control group reached an 80% confluency. Throughout the experiment, CM or DMEM was replaced every other day. The same experimental setup was followed for the cells in the negative control group treated with 10% DMSO. Cells were fixed with 150 μL 100% methanol (Merck, #Cat no: 1.06009.2511) for 20 mins at room temperature (RT). Then, formed colonies were stained with 100 μL 0.2% Crystal Violet Solution (Sigma, #Cat no: C077) for 15 mins at RT. Background staining was eliminated by washing the wells twice with 100 μL ddH_2_O. The plate was left to dry out overnight. The formed colonies were photographed under the microscope and then counted with the ImageJ 1.52v image analysis software (htpp://imagej.nih.gov/ij) “ColonyCounter” plugin.

### 2.5 Wound healing assay

CMs collected at 48^th^ h from LN229 and U87 cell lines were used to compare the effects of CM on the migration capacity of cells. LN229 and U87 cell lines were seeded with 0.5×10^5^ cells/well confluency to a 24-well plate (ThermoFisher Scientific) in a 1 mL culture medium and incubated for 24 hours. After incubation, culture mediums were replaced with 500 μL CMs or DMEM. After 24 hours, a gap was created by straight scratch using 200 μl pipette tips. Then, the cell medium was replaced to get rid of dead and de-attached cells, and finally, 0^th^, 6^th^ and 24^th^-hour images of cells were taken. ImageJ “MRI Wound Healing Tool” plugin was employed to analyze and quantify the closure rates of the created wounds.

### 2.6 Flow cytometry assay

The proliferation rate was evaluated using Click-IT^™^ EdU cell proliferation kit (ThermoFisher Scientific, #Cat no: C10425) by following the manufacturer’s protocol. Briefly, cells were seeded with 2×10^5^ density in a 1 mL medium in 12-well plates (ThermoFisher Scientific). After 24-hour incubation, the culture medium was replaced with a 600 μL CM or DMEM, and cells were incubated for an additional 24 hours. Following the incubation, 0.6 μL EdU (Component A) was added to each well with a final concentration of 10 μM, and cells were incubated for 2 hours. After incubation, cells were harvested using 0.25% Trypsin-EDTA and washed with 3 mL of 1% BSA (Capricorn, #Cat no: BSA-1s) in PBS (Gibco, #Cat no:003002) and centrifuged at 1500 rpm for 5 mins. Next, the supernatant was discarded, and the pellet was resuspended in 100 μL of Click-iT^®^ fixative (Component D). Cells were incubated for 15 mins at RT, washed with 3 mL %1 BSA in PBS, and pelleted with the same centrifugation conditions. The supernatant was removed, and cells were resuspended in 100 μL 1X Click-iT^®^ saponin-based permeabilization and wash reagent (Component E) and incubated for 15 mins at RT. Then, 500 μL of Click-iT^®^ reaction cocktail prepared according to manufacturers’ protocol with the rest of the components were added to each tube and incubated for 30 mins. Following the incubation, cells were washed with 3 mL 1% BSA in PBS and pelleted. Hoechst 33343 dye solution (Thermofisher Scientific, #Cat no: H3570) was diluted with PBS in 1:100 v/v, and 100 μL of diluted dye was added into each tube to label the viable cells. Flow cytometry analyses were performed with Navios EX (Beckman Coulter) instrument using 405 and 488 nm lasers to detect viable and EdU^+^ cells, respectively.

### 2.7 Gene expression profiling

To establish a putative link between the observed alterations at the cellular level and gene expression profile, the mRNA level of the genes listed in Table S1 (Demircan et al., 2021) were compared between treated and control groups. U87 and LN229 cells (3×10^5^) were seeded in a 6-well plate in a 2 mL medium and incubated for 24 h. Then, the culture mediums were replaced with a 1 mL CM or DMEM. Upon an additional 24-hour incubation in CM or DMEM, cells were harvested with 0.25% Trypsin-EDTA and washed with 1 mL PBS. TRIzol reagent (ThermoFisher Scientific, #Cat no: 15596026) was used to isolate total RNA according to the manufacturer’s instructions. The quality of isolated RNAs was checked on a 1% agarose gel, and RNAs were quantified with Qubit4 Fluorometer (Invitrogen, Thermofisher Scientific). cDNA synthesis was carried out with M-MuLV Reverse Transcriptase (New England Biolabs, #Cat no: M0253L), and qPCR reactions were conducted using gene-specific primers with SensiFAST SYBR No-ROX kit (Bioline, #Cat no: BIO-98005) as described previously (Sibai et al., 2019). GAPDH gene was used as the housekeeping gene for normalization.

### 2.8 Statistical analysis

Statistical analyses and graphing were performed in GraphPad Prism 5.0 software (GraphPad Software, Inc., San Diego, CA) and R language (4.3.2). The normality of data distribution was tested with the Shapiro-Wilk test. One-way analysis of variance (ANOVA) followed by Tukey’s post hoc test was employed for the normally distributed data, and Kruskal-Wallis test followed by Dunn’s post hoc test was employed as a non-parametric test for non-normal distributed data in MTT and CFA results. Pairwise comparisons in wound healing were analyzed using Wilcoxon or t-tests based on the normality of the data. Statistical significance was set at p-value<0.05.

## 3. Results and Discussion

GBM is classified as a grade IV central nervous system cancer that arises from astrocyte transformation and is manifested by highly fatal and aggressive characteristics (Hanif et al., 2017). Currently, it is an incurable disease due to the lack of an efficient treatment option (Davis, 2016). GBM progression and aggressiveness are driven by multiple factors such as genetic background (Davis, 2016), epigenetic alterations (Dong and Cui, 2019), and cross-talk between tumor and tumor microenvironment (Papaleo et al., 2017). Elucidation of these cellular and molecular contributors would be valuable to develop new therapeutic options for the better progression of GBM.

Conditioned medium has been extensively used in in-vitro studies to mimic the microenvironment of cellular niche and, therefore, dissect the role of secretome on molecular processes (Papaleo et al., 2017). It has been documented that CM derived from various cell types has substantial and multiple effects on target cells’ cellular morphology (Camerlingo et al., 2019), proliferation (Sawa-Wejksza et al., 2018), motility (Chen et al., 2014), epithelial-mesenchymal transition (Guo et al., 2017), gene expression (Bonitz et al., 2019), cell viability (Song et al., 2018), and death (Zhuang et al., 2018). There are some controversial data on the observed effects of CM treatment, most probably related to the content of released secretome, which is profoundly shaped by the origin of cells and the physiological conditions at secretion. Therefore, we first aimed to investigate the effect of CMs derived from cell lines on each other.

### 3.1 CMs derived from U87 and LN299 cells exhibited diverse effects on SVG p12 viability and clonogenicity

MTT and colony-forming assays were conducted to test the potential cytotoxic effects of CMs derived from cell lines on each other. Cell viability significantly increased in SVG p12 cells by 16% when exposed to U87 6^th^ h CM (p-value<0.001). U87 CM collected at other time points (12^th^ h, 24^th^ h, and 48^th^ h) did not significantly affect the viability of SVG p12 cells (Figure 1A). On the contrary, CMs derived from LN229 significantly decreased the viability of SVG p12 cells (p-value<0.001 for 12^th^, 24^th^, 48^th^ h CM and p-value<0.01 for 6^th^ h CM) (Figure 1A). The decreased rate was found 25% for 48^th^ h, 20% for 24^th^ h, 14% for 12^th^ h, and 7% for 6^th^ h CM. As a next step, the effect of CMs on SVG p12 cells’ colony-forming ability was investigated using CFA (Figure 1B-C). Statistical analyses revealed that CMs derived from U87 or LN229 cells did not significantly affect the colony formation capacity of SVG p12 cells (Figure 1C).

**Figure 1.**
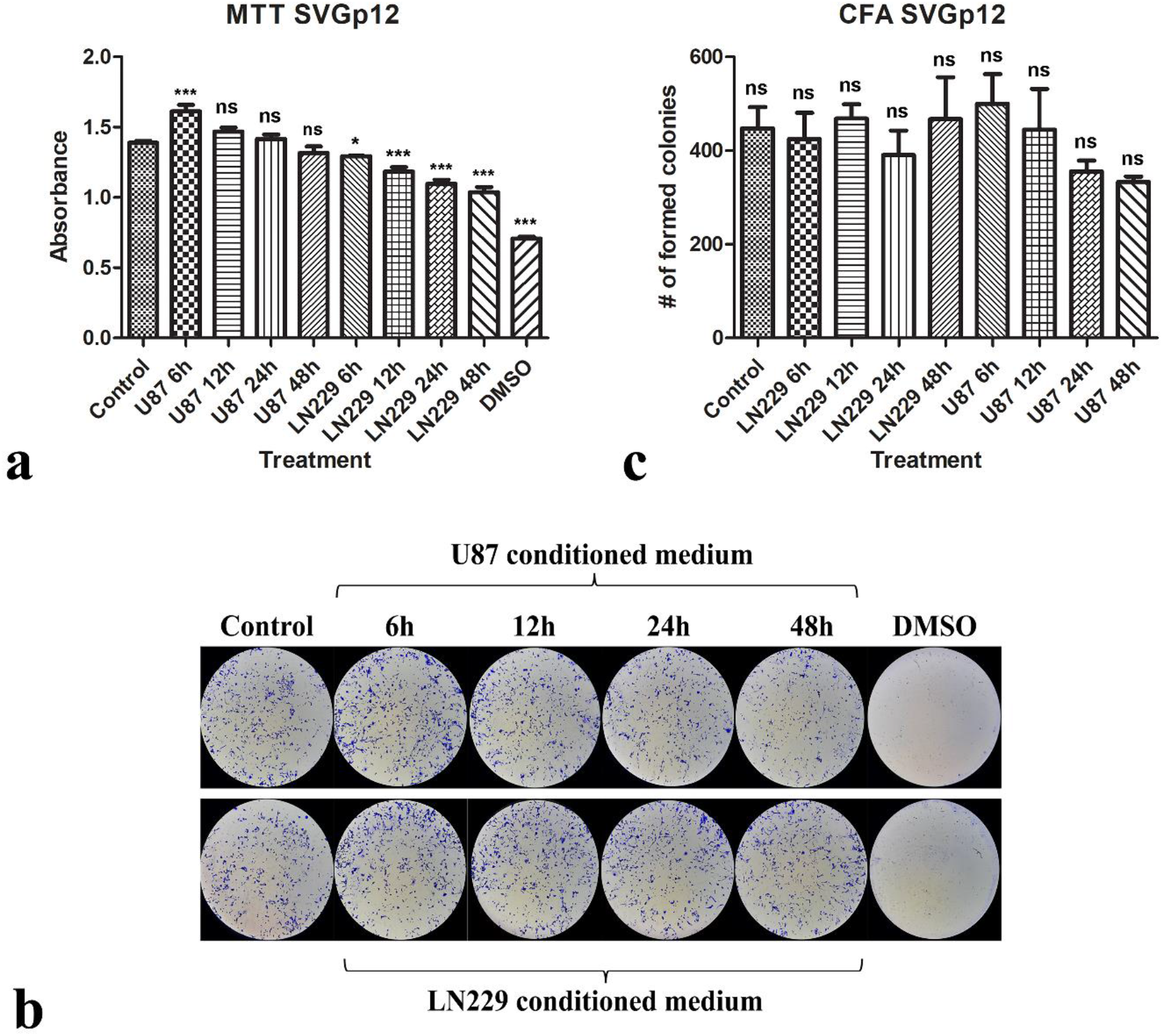
Assessing cellular viability and colony-forming ability of SVG p12 cells after CM treatments. Cellular viability and colony-forming ability were measured following the MTT and CFA protocols, respectively. a) Absorbance values obtain from SVG p12 cells by CMs of U87 and LN229, DMEM, and DMSO treatments. b) Representative images of formed SVG p12 colonies. c) Comparison of the number of formed SVG p12 colonies upon different treatments. Statistical significance was demonstrated with an asterisk. *p-value < 0.05, ***p-value < 0.001. ns; nonsignificant.

The pro-proliferative effect of U87 cells secretome content at the 6^th^ h time point was an interesting finding, and secretome at this time point might have a rich pro-proliferative signal content compared to the other time points, which should be confirmed by further studies. The observed effect may be due to the released growth factors in the 6^th^ h time point secretome. In a previous study (Guo et al., 2012), five growth factors (BDNF, VEGF, NGF, TGF-β, and NT-3) were identified in CM harvested from U87 cells. Increased proliferation rate due to released growth factors was demonstrated in other cancer types as well. Jia et al. described the increased proliferation rate of hepatocellular carcinoma (HCC) cells by the hepatocyte growth factor content of the CM collected from cancer-associated fibroblasts (Jia et al., 2013). On the other hand, CMs collected from LN229 cells at all time points reduced the viability of SVG p12 cells significantly. In a recent study, it has been demonstrated that colon cancer-derived CM repressed the monocyte proliferation and reduced the viability of THP-1 cells by arresting the cells at the G1-phase (Sawa-Wejksza et al., 2018). Another study revealed that squamous carcinoma cells viability and proliferation rate treated with mesenchymal stem cells CMs decreased significantly (Bagheri et al., 2021). Hence, CMs may have growth-limiting and anti-proliferative effects on target cells, as observed for SVG p12 cells treated with CMs obtained from LN229 cells. Notably, CMs of LN229 and U87 cells did not significantly affect the colony-forming ability of SVG p12 cells (Figure 1B, 1C). This might be due to a more naive long-term effect of CM treatment on SVG p12 cells. Alternatively, since MTT assay evaluates metabolic activity and CFA is used to test the colony formation capacity, we can speculate that CMs may decrease the metabolic activity of SVG p12 without affecting the colony-forming ability.

### 3.2 LN229 and U87 CMs collected at 48^th^ h displayed growth-limiting activity on GBM cells

In LN229 cells, the viability of cells was significantly decreased when cells were treated with CMs derived from U87 at 12^th^ (10%) 24^th^ (16%), 48^th^ (21%) h, and CM derived from SVG p12 at 48^th^ h (11%) time points (p-value<0.01 for CM harvested from U87 at 12^th^ h and p-value<0.001 for the rest) (Figure 2A). The effect of all other treatments was nonsignificant (Figure 2A). CMs collected from U87 at all time points (6^th^, 12^th^, 24^th^, 48^th^ h) significantly reduced formed LN229 colonies (p-value<0.05 for 12^th^ h treatment group; p-value<0.01 for the other treatment groups) (Figure 2B-C). 12^th^ h and 48^th^ h SVG p12 CM treatment groups exhibited a statistically significant decrease in the number of formed colonies (p-value<0.05 for 12^th^ h, and p-value<0.01 for 48^th^ h) (Figure 2C).

**Figure 2.**
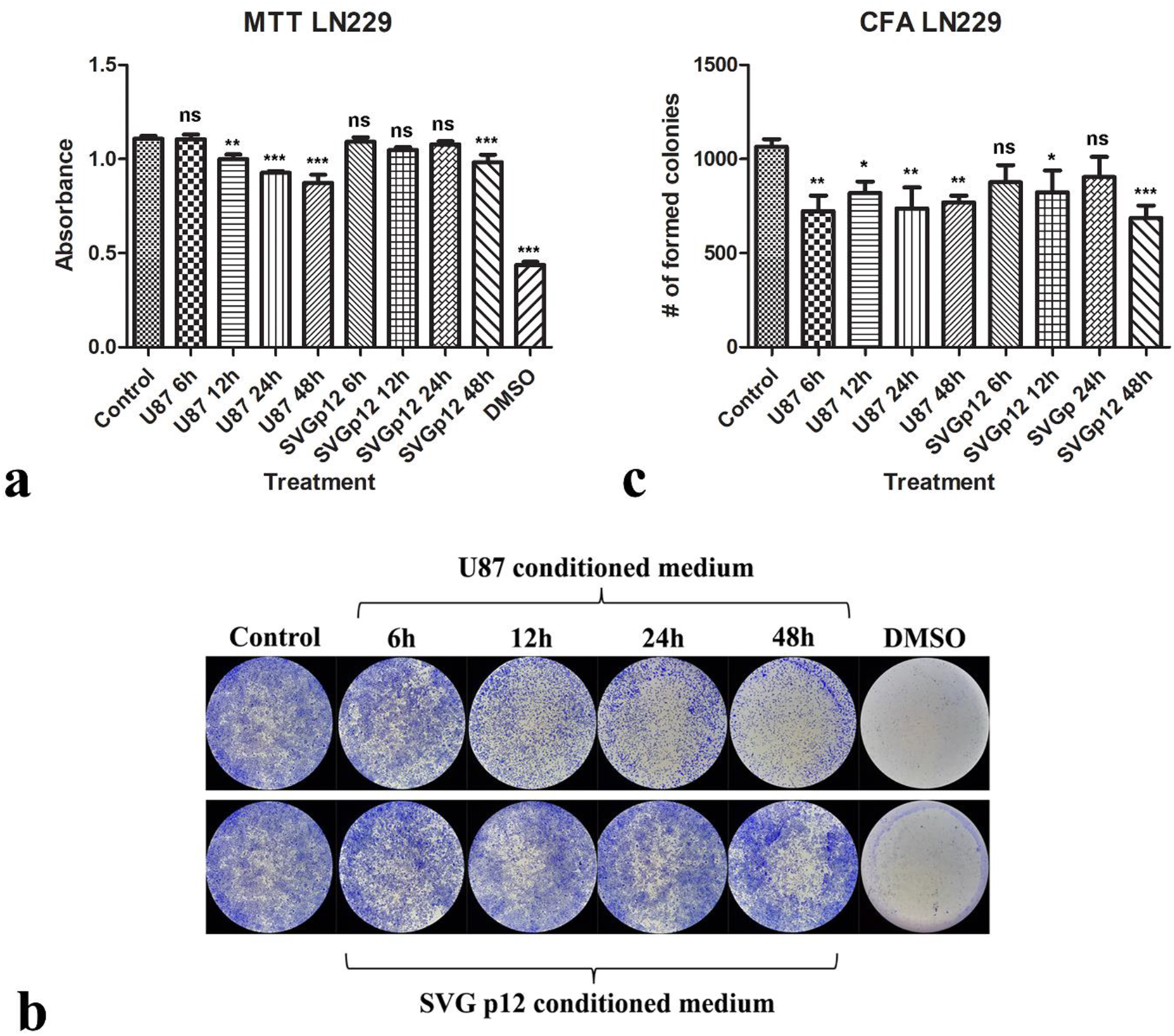
Evaluation of CMs effect on LN229 cells viability and colony formation. MTT and CFA assays were performed to evaluate the impact of CM treatment on the viability and colony formation capacity of LN229 cells. a) Absorbance levels of LN229 cells after CMs, DMEM, and DMSO treatment. b) Images of LN229 colonies formed after various treatments. c) Number of formed LN229 colonies following the treatments. Statistical significance was demonstrated with an asterisk. *p-value < 0.05, **p-value < 0.01, and ***p-value < 0.001. ns; nonsignificant.

CMs obtained from SVG p12 cells at all selected time points did not show any significant effect on U87 cells’ viability (Figure 3A) or colony-forming ability (Figure 3B-C). Based on MTT and CFA results, CMs derived from LN229 cells at 6^th^and 12^th^ h did not significantly change the viability or colony formation potential of U87 cells (Figure 3A-C). However, harvested CMs at 24^th^ and 48^th^ h significantly reduced viability of U87 cells by 11% and 13%, respectively (p-value<0.05 for 24^th^ h and p-value<0.01 for 48^th^ h treatment groups) (Figure 3A). CFA results for 24^th^ and 48^th^ h CMs of LN229 cells confirmed the MTT results by significantly decreasing the colony-forming capacity of U87 cells (p-value<0.01 for 24^th^ h and p-value<0.001 for 48^th^ h treatment groups) (Figure 3B-C).

**Figure 3.**
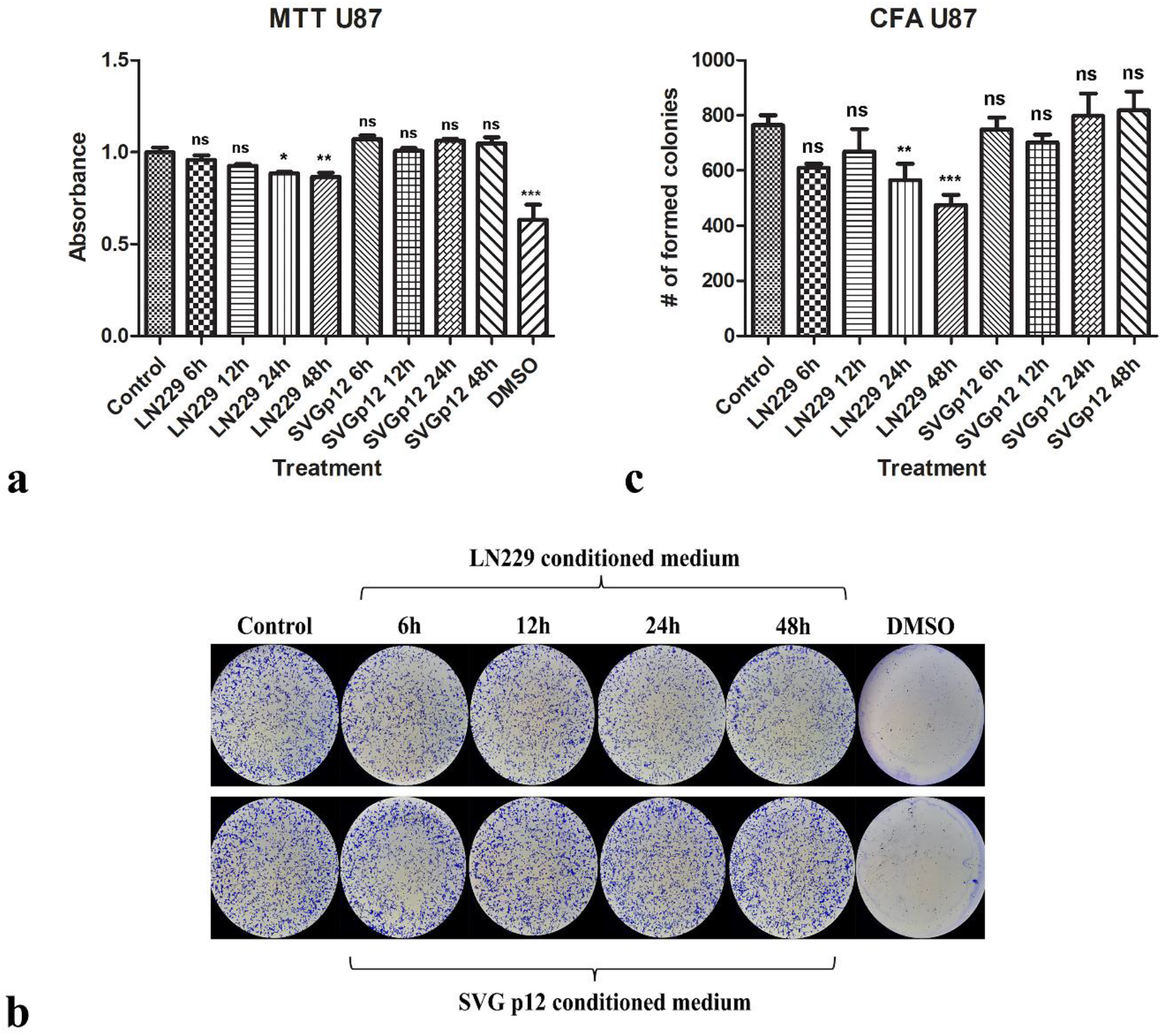
Testing CMs influence on U87 cells by MTT and CFA. Alteration of cellular viability and colony formation capacity of U87 cells. a) U87 MTT results after treatment of LN229 and SVG p12 CMs, DMEM, and DMSO. b) Representative images of U87 colonies. c) Colony numbers of U87 cells formed after LN229 and SVG p12 CMs treatment. Statistical significance is demonstrated with an asterisk. *p-value < 0.05, **p-value < 0.01, and ***p-value < 0.001. ns; nonsignificant.

SVG p12 derived CMs effect on viability and clonogenicity was limited for LN229 cells and not significant for U87 cells (Figure 2,3). In a previous study, CM obtained from astrocytes had no considerable effect on CD133^−^ GBM cells (Rath et al., 2013) which might be due to the selective uptake of the secretome components (Xu et al., 2021). U87 cells may actively use mechanisms to prevent the uptake of SVG p12 secretome elements or exclude them effectively compared to the LN229 cells. Obtained results from our MTT and CFA unveiled that GBM CMs have considerably anti-growth effects on each other, particularly for the CMs collected at the 48^th^ h time point. To explore the protein composition of 24th h CMs harvested from GBM cells (HNGC2, U87, and LN229), a proteome analysis was conducted, and proliferation related proteins such as TIMPc1, SERPIN F1, and mTOR were identified commonly as part of secretome content (Polisetty et al., 2011). Intriguingly, although CMs of GBM cell lines (U87, U373, and U251) collected after 24 h significantly increased the number of formed colonies following the reciprocal CM treatment, a considerable decrease in proliferation rate for these cells was noted (Motaln et al., 2015). Therefore, the content of secretome may act on diverse biological processes of target cells. Based on the literature, as far as we know, 48^th^ h CM of GBM cells has not been studied extensively before. It is plausible to suggest that 48^th^ h CM of GBM cells could contain anti-proliferative signals along with pro-proliferative signals, which should be addressed in future studies.

Considering our MTT and CFA results which highlighted that LN229 and U87 48^th^ h CMs have a significant growth-limiting effect on each other, we performed the subsequent experiments using 48^th^ h CMs. Hereafter, “LN229-CM” and “U87-CM” will be used to refer to CM harvested at 48^th^ h from LN229 cells and U87 cells, respectively.

### 3.3 LN229-CM significantly decreased the migration capacity of U87 cells

Wound healing assay was conducted to evaluate the effect of CMs on the migration capacity of GBM cells. First, we performed a statistical analysis within each group. The wound area was almost completely closed 24 h post scratch for U87 cells cultured in DMEM (p-value<0.001) (Figure 4A-B). The size of the scratch at the 6^th^ h time point was significantly reduced for U87 cells in DMEM compared to the 0^th^ h scratch size (p-value<0.001) (Figure 4B). The decreasing trend in wound size was extended to 24^th^ h compared to the 6^th^ h time point in control U87 cells (p-value<0.001) (Figure 4B). Interestingly, there was no significant difference in scratch size between 0^th^ h and 6^th^ h for LN229-CM treated U87 cells (Figure 4B).

**Figure 4.**
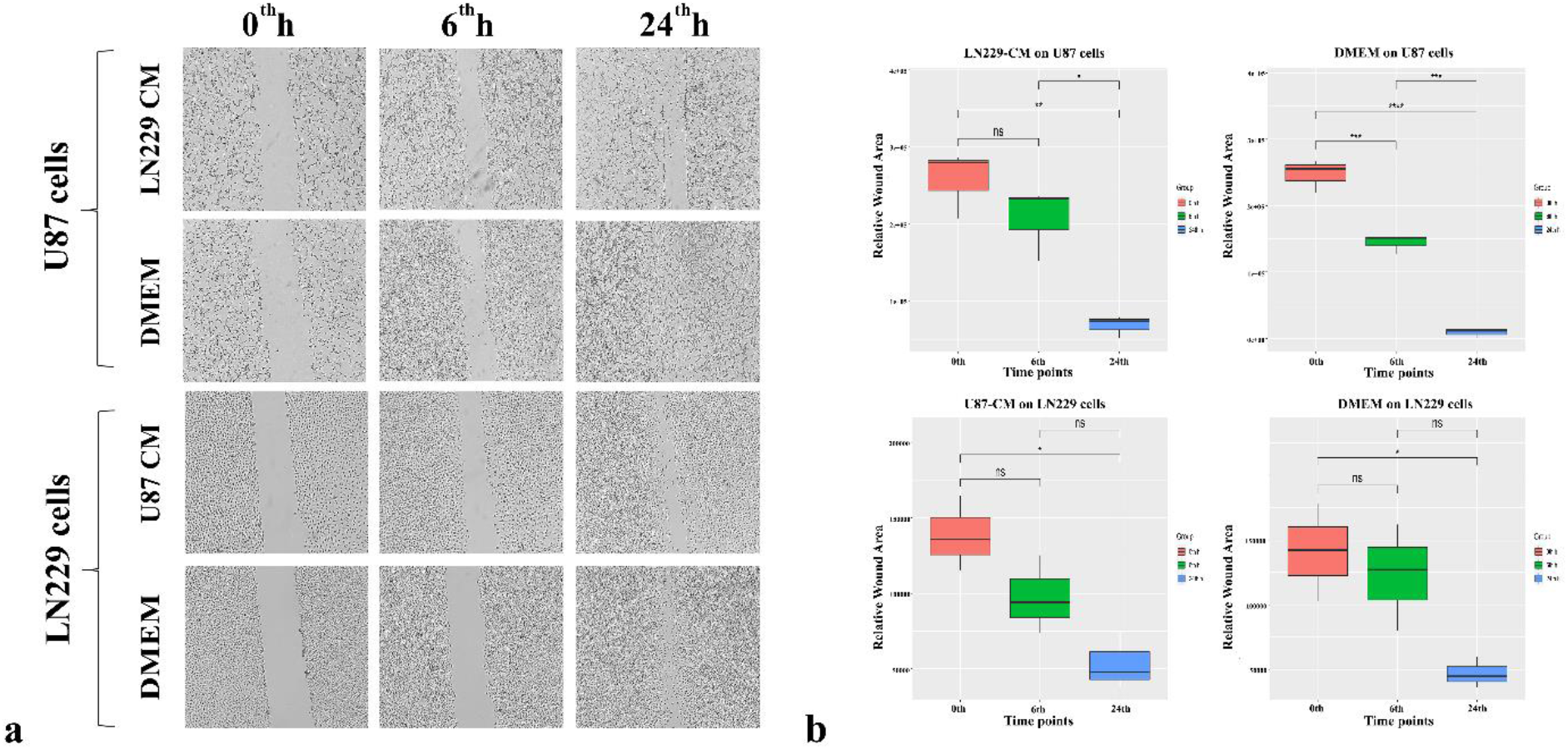
Effect of the CMs on migration capacity. Wound healing assay was employed to test the migration capacity of GBM cells when treated with CMs or DMEM. The effect of CMs harvested from U87 and LN229 at 48^th^ h was examined. Representative images of U87 and LN229 cells at 0^th^, 6^th^ and 24^th^ h post scratch were shown. Statistically significant groups are displayed with an asterisk. *p-value < 0.05, **p-value < 0.01, and ***p-value < 0.001. ns; nonsignificant. U87-CM; 48^th^ h CM of U87 and LN229-CM; 48^th^ h CM of LN229 cells.

On the other hand, the closure rate was found significant between 24^th^ h compared to 0^th^ h and 6^th^ h time points (p-value<0.01 and p-value<0.05, respectively) (Figure 4B).

For control and U87-CM treated LN229 cells, the 6^th^ h point wound closure rate was nonsignificant. Control LN229 cells significantly closed the wound at the 24^th^ h time point in comparison to the 0^th^ h time point (p-value<0.05) Figure 4B). Wound area between 6th h and 24th h time points was also compared, and the change in wound size between these time points was found nonsignificant for control LN229 cells (Figure 4B). Wound size at 24^th^ h time point for U87-CM treated LN229 cells was significantly decreased compared to 0^th^ h time point (p-value<0.05) (Figure 4B). For the same treatment, the comparison of scratch size between 24th h and 6th h time points was nonsignificant (Figure 4B).

Next, we compared the success of healing between treated and control groups for the same cell line. There was a significant difference in wound healing rate for both 6th h and 24th h time points between LN229-CM treated and non-treated U87 cells (p-value<0.05) (Table S4). For LN229 cells, there was no significant difference between CM treatment and control groups (Table S4).

To summarize the obtained results from the wound healing assay, a significant anti-migratory effect of LN229-CM on U87 cells was observed; however, U87-CM did not significantly affect the LN229 migration ability. Decreased migration potential of a variety of cells following the CM treatment has been documented. For example, CM collected from the B16F10 mouse melanoma cancer cells reduced the macrophage motility (Go et al., 2013). In another study, adipose-derived stem cell CM inhibited the migration of B16 melanoma cells (Lee et al., 2015). On the other hand, there are some contrary reports on the migratory effect of CM. CMs derived from malignant breast cancer cell lines MDA-MB-231 and MDA-MB-453 increased the migration capacity of non-tumorigenic MCF10A cells (Guo et al., 2017). In accordance with this, osteoblast CM increased migration of the MCF-7 breast cancer cell line (Chen et al., 2014). Therefore, we believe that the diverse and even opposite observed effect of CM on the migration capacity might be related to the type of cells used to collect the CM and the target cells treated by the collected CM. More studies to unveil the content of LN229 and U87 CMs are necessary to extend our current understanding of the contribution of CMs to cell migration capacity.

### 3.4 Anti-proliferative effect of U87-CM on LN229 cells was detected

From the MTT and CFA results, we speculated that the growth-limiting effect of LN229-CM and U87-CM might be linked to altered proliferation rate. To further interrogate the anti- or pro-proliferative impact of CMs, EdU based proliferation assay was carried out. For U87-CM treated LN229 cells, a significant decrease in proliferation rate compared to non-treated LN229 cells was observed (p-value<0.001) (Figure 5A). However, LN229-CM treatment did not significantly affect the proliferation rate of U87 cells (Figure 5B).

**Figure 5.**
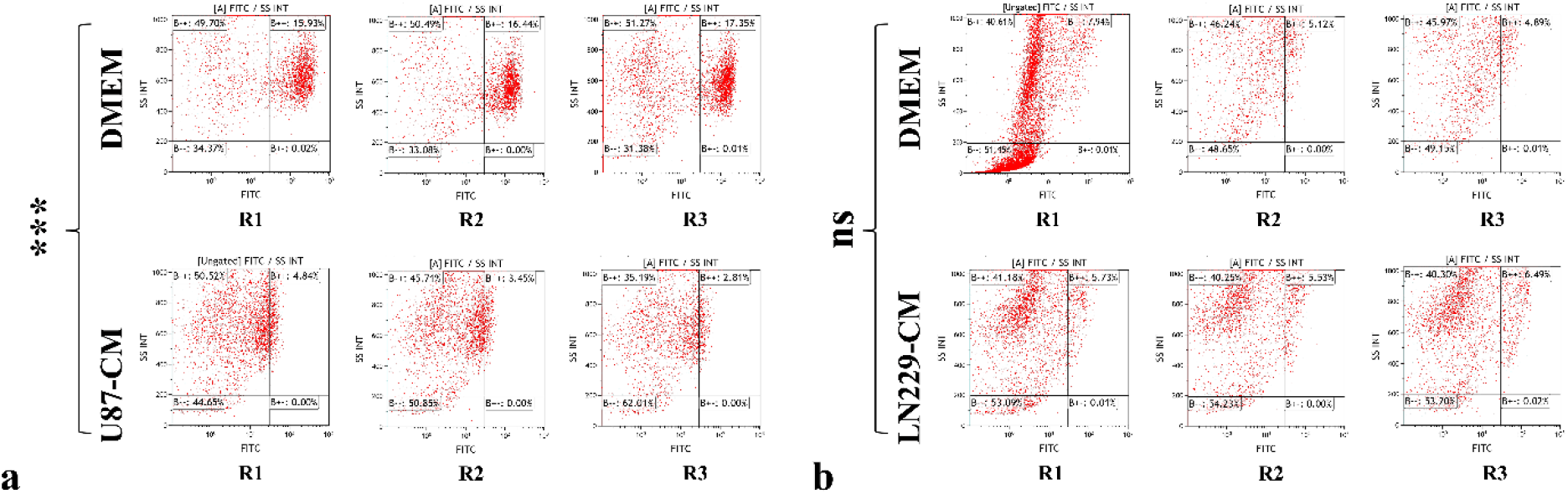
Evaluation of CMs’ impact on proliferation rate. EdU assay was applied to assess the outcome of CM treatment on proliferation rate. EdU incorporation in a) CM harvested from U87 cells at 48^th^ h and DMEM treated LN229 cells, and b) CM harvested from LN229 cells at 48^th^ h and DMEM treated U87 cells. FITC^+^ cells are considered as positive EdU incorporating cells. ***p-value < 0.001. ns; nonsignificant, R; replicate. U87-CM; 48^th^ h CM of U87 and LN229-CM; 48^th^ h CM of LN229 cells.

Cell proliferation results confirmed that U87-CM contains anti-proliferative factors for LN229 cells. This may explain the MTT and CFA results where we observed the growth-limiting activity of U87-CM on LN229 cells. In contrast, no significant effect of LN229-CM on the proliferation rate of U87 cells highlights the necessity of more studies to explore the source of LN229-CM growth-limiting factors, which remained unclear. Evaluation of the cell death rate of U87 cells upon LN229-CM treatment might be helpful to deal with this question. As shown in a previous study, CMs harvested from GBM cells exhibited potential inhibitory effects on each other (Motaln et al., 2015). In that study, it has been reported that the CM of U251 and U373 cells decreased the proliferation rate of U87. Interestingly, CM of U87 cells did not significantly affect the proliferation rate of U251 or U373 cells pinpointing the diverse impact of GBM CMs on each other as observed in our study.

### 3.5 Levels of both pro-and anti-proliferative genes in U87 and LN229 cells were altered following the CM treatment

To expand the molecular basis of the alterations in cellular characteristics upon CM treatment, control, and treated cells’ gene expression profile was investigated and compared. For both LN229-CM treated U87 cells and U87-CM treated LN229 cells, the gene expression profile was changed considerably for the genes summarised in Table S1. Relative expression levels for different groups are demonstrated in Table 1.

**Table 1.** Gene Expression Profiles of GBM cells after treatment by CMs.

| LN229 cells |  |  |  | U87 cells |  |  |  |
| --- | --- | --- | --- | --- | --- | --- | --- |
| Gene | Treatment Group | Expression | Significance | Gene | Treatment Group | Expression | Significance |
| MKI67 | U87-CM vs. control | ↑ | * | MKI67 | LN229-CM vs. control | ↑ | ** |
| PCNA | U87-CM vs. control | ↑ | * | PCNA | LN229-CM vs control | ↑ | *** |
| CCNE1 | U87-CM vs. control | ↑ | * | CCNE1 | LN229-CM vs. control | ↑ | *** |
| BIRC5 | U87-CM vs. control | ↔ | ns | BIRC5 | LN229-CM vs. control | ↑ | *** |
| AKT | U87-CM vs. control | ↓ | ** | AKT | LN229-CM vs. control | ↑ | ** |
| BCL2 | U87-CM vs. control | ↔ | ns | BCL2 | LN229-CM vs. control | ↔ | ns |
| CDK4 | U87-CM vs. control | ↓ | *** | CDK4 | LN229-CM vs. control | ↔ | ns |
| CDK6 | U87-CM vs. control | ↔ | ns | CDK6 | LN229-CM vs. control | ↔ | ns |
| CCND1 | U87-CM vs. control | ↑ | *** | CCND1 | LN229-CM vs. control | ↔ | ns |
| DNMT1 | U87-CM vs. control | ↔ | ns | DNMT1 | LN229-CM vs. control | ↔ | ns |
| GADD45A | U87-CM vs. control | ↑ | * | GADD45A | LN229-CM vs. control | ↔ | ns |
| CDKN1A | U87-CM vs. control | ↔ | ns | CDKN1A | LN229-CM vs control | ↔ | ns |
| CDKN1B | U87-CM vs. control | ↑ | *** | CDKN1B | LN229-CM vs. control | ↔ | ns |
| ATM | U87-CM vs. control | ↑ | * | ATM | LN229-CM vs. control | ↔ | ns |
| PUMA | U87-CM vs. control | ↑ | *** | PUMA | LN229-CM vs. control | ↔ | ns |
| BAX | U87-CM vs. control | ↑ | * | BAX | LN229-CM vs. control | ↑ | * |
| TP53 | U87-CM vs. control | ↔ | ns | TP53 | LN229-CM vs. control | ↑ | ** |
| BRCA1 | U87-CM vs. control | ↑ | ** | BRCA1 | LN229-CM vs. control | ↑ | *** |

Among the pro-proliferative genes, the expression levels of the following ones were evaluated. Mammalian Ki-67 protein (MKI67) is a nuclear protein associated with cellular proliferation, and its expression level is widely used as a diagnostic and prognostic cancer marker (Menon et al., 2019). Proliferating cell nuclear antigen (PCNA) plays an essential role in DNA replication and repair mechanisms (Boehm et al., 2016). Cyclin-dependent kinases (CDKs) regulate the cell cycle and coordinate the cycle progression (Otto and Sicinski, 2017). CDK encoding genes CDK4 and CDK6 form a complex to facilitate the G1/S transition of the cell cycle. Cyclin D1 (CCND1) is the regulatory component of the CDK4/6 complex, and activation of the CDK4/6 complex by binding of CCND1 negatively regulates the Rb protein. Inactivation of Rb results in the release of the E2F complex to stimulate the expression of S-phase-specific genes (Ding et al., 2020). CCNE1 gene encodes cyclin E1 protein, which forms a complex with CDK2 kinase to regulate the G1/S transition (Hwang and Clurman, 2005). Overexpression of these genes has been detected in various cancers and found to be associated with chemoresistance of GBM (Liang et al., 2019) and epithelial ovarian cancer (Gorski et al., 2020), metastasis in gastric carcinoma (Kim et al., 2019), and poor prognosis in breast cancer (Zhao et al., 2019). BCL-2, as a potent inhibitor of pro-apoptotic proteins, forms channel complexes on mitochondria membranes to release the mitochondrial components (Shamas-Din et al., 2013). Survivin, BIRC5 encoded protein, is a member of the inhibitor of apoptosis protein (IAP) family that inhibits the activity of the caspases (Jaiswal et al., 2015). Hence, high survivin expression inhibits apoptosis in cancer cells and is associated with poor prognosis and overall survival (Kelly et al., 2011). AKT protein is a serine-threonine kinase and regulates various cellular functions, such as proliferation, survival, and angiogenesis (Yu and Cui, 2016). DNA-methyltransferases 1 enzyme catalyzes de-novo DNA methylation (Hervouet et al., 2018). Elevated levels of DNMT1 were shown in astrocytic tumor specimens, and its linkage to the high-grade tumor samples was found to be correlated with O^6^-methylguanine-DNA methyltransferases (MGMT) expression (Rahman et al., 2015).

In U87-CM treated LN229 cells, MKI67, PCNA, CCNE1, and CCND1 were significantly upregulated, and CDK4 and AKT were significantly downregulated compared to the non-treated control. No significant difference in expression levels of BCL2, CDK6, BIRC5, and DNMT1 was detected between CM treated and non-treated U87 cells. We can conclude that CM treatment stimulates a mosaic gene expression pattern for pro-proliferative genes based on the obtained results. Although many pro-proliferative genes were found as significantly upregulated upon CM treatment, a significantly diminished level of CDK4 was noteworthy. As in many other cancer cells, CDK4/6 inhibition was shown to inhibit the cellular proliferation of GBM cells (Cao et al., 2020). Decreased proliferation rate and viability of U87-CM treated LN229 cells might be connected with the downregulation of the CDK4 gene. Inadequate formation of CDK4/6-Cyclin D1 complex due to a decrease in CDK4 level may account for the observed decline in viability following the CM treatment (Figure 2A-C). Additionally, non-elevated levels of anti-apoptotic factors may have further contributed to reducing viability.

On the other hand, in LN229-CM treated U87 cells, increased MKI67, PCNA, CCNE1, BIRC5, and AKT levels were detected. For the BCL2, CDK6, CDK4, DNMT1, and CCND1 genes, no significant difference in expression levels was found between CM treated and non-treated U87 cells. Upregulated levels of pro-proliferative genes in U87 cells following the CM treatment do not agree with the MTT and EdU results. The proliferation rate of U87 cells was similar between CM treated and control cells. Moreover, based on the MTT assay, a significant negative effect of CM on U87 cells viability was observed. The dual impact of CM to activate both the proliferation and cell death mechanisms might explain this contradiction.

To get more insights into the anti-proliferative activity of CMs, we evaluated the expression level of several anti-proliferative genes. CDKN1A gene encodes cyclin-dependent kinase inhibitor 1A protein, also known as p21, and acts as an anti-proliferative regulatory protein (Abbas and Dutta, 2009). CDKN1B encoded protein, p27, binds to CDK2-cyclin E1 or CDK4-cyclin D1 and prevents their activation (Lim and Kaldis, 2013). GADD45A is a cellular-senescence regulatory gene associated with S-phase entry and induces growth arrest (Tamura et al., 2012). As shown in a previous report, cell-cycle arrest and DNA repair roles of GADD45A in a p53-dependent manner inhibit the proliferation and growth of bladder cancer cells (Han et al., 2019). ATM gene, encoding a vital cell cycle checkpoint kinase, regulates the p53 and BRCA1 in response to DNA damage (Stracker et al., 2013). Roles in homology-directed repair and mismatch-repair mechanisms position the BRCA1 as an essential tumor-suppressor gene (Chen et al., 2018). Tp53 is a nuclear transcription factor that regulates cellular stress response by activating genes involved in cell cycle arrest, DNA repair, and pro-apoptotic genes (Chen, 2016). Under normal conditions, post-translational modifications inhibit the protein’s activation and nuclear translocation (Chen, 2016). A pro-apoptotic factor, BAX, is an organelle membrane-associated protein that can induce an opening on the mitochondria via a voltage-dependent ion channel or by forming an oligomeric pore (Kilbride and Prehn, 2013). PUMA is a pro-apoptotic factor that interacts with antiapoptotic Bcl-2 family members to release the Bax or Bak to induce pores’ formation on mitochondria (Hatok and Racay, 2016).

The expression level of GADD45A, CDKN1B, BAX, ATM, BRCA1, and PUMA genes was revealed as significantly upregulated in CM treated LN229 cells compared to the control group. Upon CM treatment, no significant difference at mRNA level for CDKN1A and TP53 genes was determined. Increased anti-proliferative activity of genes in U87 CM treated LN229 cells provides plausible explanations on the decreased viability (Figure 2A-C) and proliferation rate (Figure 5A). Lower EdU positive cells after CM treatment may be linked to the inhibition of cell cycle progression by the increased GADD45A and CDKN1B levels. Moreover, upregulated pro-apoptotic factors may account for the increased rate of cell death, which may further decrease the number of alive cells following the CM treatment. Follow-up studies to evaluate the apoptosis rate should support this prior result.

In contrast, a similar expression level for CM treated and non-treated U87 cells was observed for most anti-proliferative genes. There was no significant difference in GADD45A, CDKN1A, CDKN1B, ATM, and PUMA levels between treated and control groups, and BAX, BRCA1, and TP53 were significantly upregulated in CM treated cells. Elevated levels of BAX and TP53 highlight the anti-proliferative content of harvested CM.

Altogether, CM treatment activates the expression of pro-and anti-proliferation-related genes, which introduces a possible confound in matching the proliferative effect of CMs with previous results indicating a negative regulatory function of CMs on viability. Decreased viability upon CM treatment might be associated with an elevated apoptosis and cell cycle arrest rate, which should be addressed in future studies.

## 4. Conclusion

Here, in this study, we used CMs of GBM cell lines on each other to investigate the effect of CMs at the cellular and molecular levels. Cytotoxicity, clonogenicity, migration capacity, proliferation rate, and gene expression profiles of cells treated with CMs were compared with non-treated cells. We demonstrated that CMs harvested at 48^th^ h decreased the viability and migration and modulated the gene expression of target cells. Activation of pro-and anti-proliferative gene expression programs underlines the necessity of further studies to dissect the molecular content of secretome. It remains an open question whether the apoptosis and cell-cycle arrest are associated with the decreased viability of the CM-treated cells. Another critical question to study is the potential effects of CMs obtained from the different grade glioma cells. Future studies on glioma cell lines at different pathological grades using cutting-edge proteomics technologies would provide new insights into the molecular signature of CM and its effect on target cells.

## Abbreviations

GBM =: Glioblastoma Multiforme
CM =: Conditioned Medium
U87-CM =: 48^th^ h conditioned medium of U87
LN229-CM =: 48^th^ h conditioned medium of LN229

## Authors’ Contributions

Turan Demircan: Conceptualization, Investigation, Methodology, Project administration, Original draft preparation, Reviewing and Editing. Mervenur Yavuz: Formal analysis, Investigation, Validation, Visualization, Original draft preparation, and Editing. Siddika Akgül: Investigation, Validation, Visualization. Egemen Kaya: Investigation. All authors have read and approved the final version for publication.

## Acknowledgment

The authors would like to thank Demircan Lab members for their helpful comments on the manuscript.

